# Study of pre-synaptic internalisation in human schizophrenia brains

**DOI:** 10.1101/2020.05.22.110759

**Authors:** Makis Tzioras, Anna J. Stevenson, Delphine Boche, Tara L. Spires-Jones

## Abstract

**Aims:** Efficient synaptic communication is crucial to maintain healthy behavioural and cognitive processes. Individuals affected by schizophrenia present behavioural symptoms and alterations in decision-making, suggesting altered synaptic integrity as the support of the illness. It is currently unknown how this synaptic change is mediated in schizophrenia, but microglia have been proposed to act as the culprit, actively removing synapses pathologically. Here, we aimed to explore the interaction between microglia and synaptic uptake in human post-mortem tissue.

**Methods:** We assessed microglial activation and synaptic internalisation by microglia in a post-mortem human tissue of 10 control and 10 schizophrenia cases. Immunohistochemistry was performed to identify microglia (Iba1 and CD68) and the presynaptic terminals (synapsin I).

**Results:** We found no difference in microglial expression, nor a difference in pre-synaptic protein level phagocyted by microglia between the two groups.

**Conclusions:** Our findings are consistent with the brain imaging studies in schizophrenia implying that microglia play a role mainly during the early phases of the disease, by example in active synapse remodelling, which is not detected in the chronic stage of the illness.

## Introduction

Efficient synaptic communication is crucial to maintain healthy behavioural and cognitive processes. In neurodevelopmental diseases, like schizophrenia, affected individuals can exhibit behavioural symptoms like psychosis, hallucinations and alterations in decision-making. A reduction in cortical grey matter volume and enlarged ventricles in the brains of schizophrenia cases has been consistently reported [1,2]. This reduction in cortical volume is likely to be an outcome of neuronal and synaptic loss, which has also been reported in schizophrenia but the results have varied between brain area and synaptic markers examined [3–7]. A meta-analysis of the expression of synaptic markers in the disease has shown reduced levels of pre-synaptic markers, including synaptophysin and synapsin, in the hippocampus and frontal cortex which are heavily implicated in schizophrenia, but not in unaffected areas like the temporal and occipital lobes [8]. Synapses are crucial mediators of brain communication [9–11], and so, such synaptic alterations can have an impact on brain network connectivity, a process known to be affected in schizophrenia [12]. There are several factors during brain development that influence brain connectivity, with non-neuronal contributors playing an important role in synaptic formation and network maturation [13,14]. One of these non-neuronal contributors are microglia, the resident brain immune cells and primary phagocytes of the brain [14–17], which have been shown to facilitate neural network shaping in development by phagocytosing synapses using the complement system [18–20]. However, microglia can be aberrantly involved in synaptic elimination in non-physiological contexts, like observed in animal models of Alzheimer’s disease [21,22]. In schizophrenia, few experimental studies have explored the role of microglia in synaptic loss with evidence to suggest their involvement in excessive synaptic pruning [23]. Whether this is true in the human brain in schizophrenia is unknown. Here, we perform a human post-mortem study to investigate the role of microglia in synaptic engulfment in schizophrenia.

## Methods

### Human tissue

Ten cases with a confirmed diagnosis of schizophrenia and 10 non-neurological and non-neuropathological controls were obtained from the Corsellis Collection (Table 1). Dorsal prefrontal cortex, an area showing neuroimaging abnormalities with reduction of the grey matter volume in chronic schizophrenia [1], was investigated for all cases. Cases with any other significant brain pathologies such as infarct, tumour, or traumatic brain injury were excluded from the study. Controls with no history of neurological or psychiatric disease or symptoms of cognitive impairment were matched to cases as possible. No difference in age at death and in post-mortem delay was detected between the 2 groups. To minimize the time in formalin, which has an effect on the quality of the immunostaining, the selection was performed on the availability of formalin-fixed paraffin embedded tissue, and thus on blocks processed at the time of the original post-mortem examination. Characteristics of the cohorts are provided in Table 1.

**Table 1:** Demographic, clinical and *post-mortem* characteristics of control and schizophrenia cases

| Cases | Ctrl (n=10) | Sz (n=10) | P value |
| --- | --- | --- | --- |
| Sex | 4F:6M | 2F:8M |  |
| Age at death (years, mean ± SD) | 64.40 ± 19.78 | 64.80 ± 20.37 | P = 0.94 |
| Post-mortem delay (hours, mean ± SD) | 61.90 ± 51.23 | 50.60 ± 24.52 | P = 0.61 |
| Age of onset (years, mean ± SD) | NA | 36.50 ± 13.81 | |
| Duration of illness<br>(years, mean ± SD) | NA | 35.13 ± 21.85 | |
| Cause of death |  |  |  |
| Cardiovascular disease | 8 | 5 |  |
| infection/inflammation | 1 | 3 |  |
| Trauma | 1 | 1 |  |
| Others* | 0 | 1 |  |

Ctrl, neurologically/cognitively normal controls; Sz, Schizophrenia cases; F, female, M, male; SD, standard deviation; NA, Not Applicable; \*foreign body in respiratory tract

### Immunohistochemistry

Paraffin-embedded tissue was cut at 7μm thickness on a microtome and mounted on Superfrost glass slides. The tissue was dewaxed in xylene, followed by rehydration in 100% EtOH, 90% EtOH, 70% EtOH, 50% EtOH, and finally water for 3 minutes each. Citric acid pH6 (VectorLabs, H-3300) was used for heat-mediated antigen retrieval by pressure cooking for 3 minutes at the steam setting. The slides were incubated for 5 minutes with autofluorescence eliminator reagent (Merck Millipore, 2160) to reduce background, followed by another 5 minutes incubation with Vector TrueView autofluorescence quenching kit (VectorLabs, SP-8400) to reduce red blood cell autofluorescence. Sections were blocked in 10% normal donkey serum (Sigma Aldrich, D9663-10ML) and 0.3% Triton X-100 (T8787-100ML) for 1 hour at room temperature. Microglia were stained with Iba1 (Abcam, ab5079, goat polyclonal, 1:500), and CD68 (DAKO, M0876, mouse monoclonal, 1:100), and pre-synaptic terminals with synapsin I (Sigma Aldrich, AB1543P, rabbit polyclonal, 1:750), overnight at 4°C in a humid chamber. All primary antibodies were diluted in the blocking solution described above. The following cross-adsorbed secondary antibodies were used: donkey anti-goat A647 (Thermo Fisher Scientific, A32849), donkey anti-mouse A594 (Thermo Fisher Scientific, A32744), and donkey anti-rabbit A488 (Thermo Fisher Scientific, A32790). All secondary antibodies were diluted in phosphate buffer saline (PBS) (Thermo Fisher Scientific, 70011036). For tissue washes, 10X PBS was diluted in water to 1X concentration, with the addition of 0.3% Triton X-100 in washes prior to primary antibody incubation. Nuclei were counterstained with DAPI (1μg/ml) (D9542-10MG, Sigma-Aldrich).

### Confocal microscopy and image analysis

Twenty images were taken randomly throughout all cortical layers of the grey matter for each case using a Leica TCS8 confocal microscope with a 63x oil immersion objective. Acquisition parameters were kept constant for all images and cases. Lif files were split into tiff files, and batch analysed on ImageJ (version 1.52p, Wayne Rasband, NIH, USA) using a custom co-localisation and thresholding macro. Images from different cases were also manually analysed to ensure the macro was accurately detecting positive signal and excluding background. For synaptic internalisation by microglia we chose to analyse the colocalization of CD68 with Syn1, and also normalised to either CD68 or Iba1 burden. Data are expressed as protein burden (%) defined as the area fraction of each image labelled by the antibody. 3D reconstruction images were generated in ParaView 5.8.0. All experiments and analyses were blinded to the experimenter.

### Ethics

Ethical approval was provided by BRAIN UK, a virtual brain bank which encompasses the archives of neuropathology departments in the UK and the Corsellis Collection, ethics reference 14/SC/0098. The study was registered under the Ethics and Research Governance (ERGO) of the Southampton University (Reference 19791).

### Statistics

R Studio version 3.6.0 (2019-04-26) was used for statistical analysis [24]. Linear mixed-effects models were used to examine the effect of disease status on microglial burdens and CD68-Synapsin I co-localisation. This test was chosen because it allows all 20 images taken per case to be considered while accounting for non-independence, instead of a single mean value per case, allowing for a more powerful analysis on the results. QQ plots were generated in R Studio to check the residuals were normally distributed, which is an assumption of the mixed-effects model. To meet the assumptions of the test, all datasets were Tukey transformed prior to analysis (untransformed data presented in graphs). GraphPad Prism 8 was used for generating bar graphs with a mean value plotted per case, represented as a dot. We considered p ≤0.05 as significant.

## Results

### Microgliosis

We studied post-mortem brains from 10 control (mean age 64.40 ± 19.78) and 10 confirmed schizophrenia cases (mean age 64.80 ± 20.37) from dorsolateral pre-frontal cortex (DLPFC, Brodmann area 46) which is affected in schizophrenia [1]. Gliosis is commonly observed during loss of brain homeostasis. We examined microglial burden using Iba1 which labels the microglial cytoplasm and reflect microglial motility and homeostasis Iba1 is considered as a pan-microglial marker and has been described increased in a subset of neurodegenerative diseases [25]. The other microglial marker, CD68, labels the lysosomal compartment of microglia revealing phagocytosis [26] (Figure 1A-B), and is increased in many neurological diseases, including Alzheimer’s disease [27] and stroke [28]. By thresholding for each of the markers and quantifying their respective burdens, we found there was no difference in either Iba1 (p=0.315) or CD68 (p=0.794) burdens between the schizophrenia and control cohorts (Figure 1) (full statistical outcomes found in Supplementary Table 1). Furthermore, there was no difference in the co-localisation between CD68 and Iba1 in controls and schizophrenia brains (p=0.639), suggesting the co-expression of the two markers per single cell is unchanged (Figure 1E).

**Figure 1.**
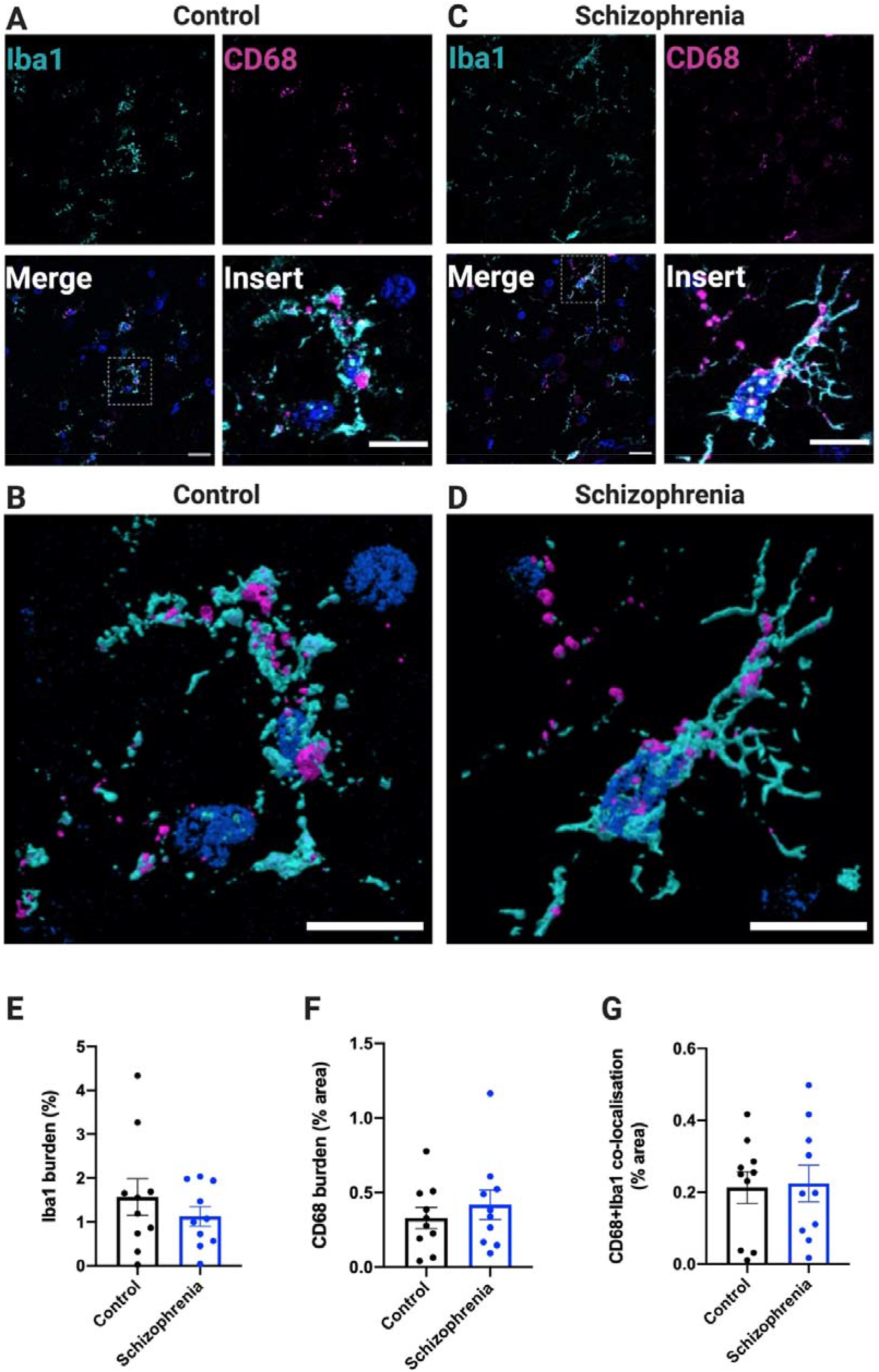
Microgliosis burdens unchanged in control and schizophrenia tissue. Representative confocal images of immunohistochemistry stained sections for the microglial markers Iba1 (cyan) and CD68 (magenta) in control (A and B) and schizophrenia (C and D) tissue. Nuclei are counterstained with DAPI. Scale bars in large images represent 20μm and 10μm in the expanded inserts (denoted by dotted white lines). The insert images of A and B are represented as 3D-reconstruction in B and D, respectively (scale bar, 10μm). 3-D reconstructions made on ParaView. Quantification of Iba1 burdens (% area), CD68 burdens (% area) and Iba1+CD68 co-expression (% area) are shown in panels E, F, and G, respectively. Each data point represents a mean of 20 images taken per case, where n=10 per group. Linear mixed-effects model showed no difference in microgliosis between the control and schizophrenia cases in the above comparisons at a significance threshold of p≤0.05.

### Synaptic engulfment

Though no difference in microglial burdens between the two cohorts was observed, we aimed to assess whether microglia were involved in synaptic engulfment in schizophrenia. To do this, we quantified the amount of co-localisation between synapsin I and CD68 (% area), as a measure of engulfed synaptic material in the microglial phago-lysosomal compartment (Figure 2). We found no difference in synaptic engulfment by microglia between the schizophrenia and control cases (p=0.413) (Figure 2). Furthermore, when we normalised this co-localisation to their respective CD68 or Iba1 burdens, there was still no statistical difference between schizophrenia and control tissue (p=0.167 and p=0.964 respectively) (Figure 2). Our data therefore suggest that at the time of death, microglia do not appear to be involved in aberrant synaptic internalisation in patients with schizophrenia.

**Figure 2.**
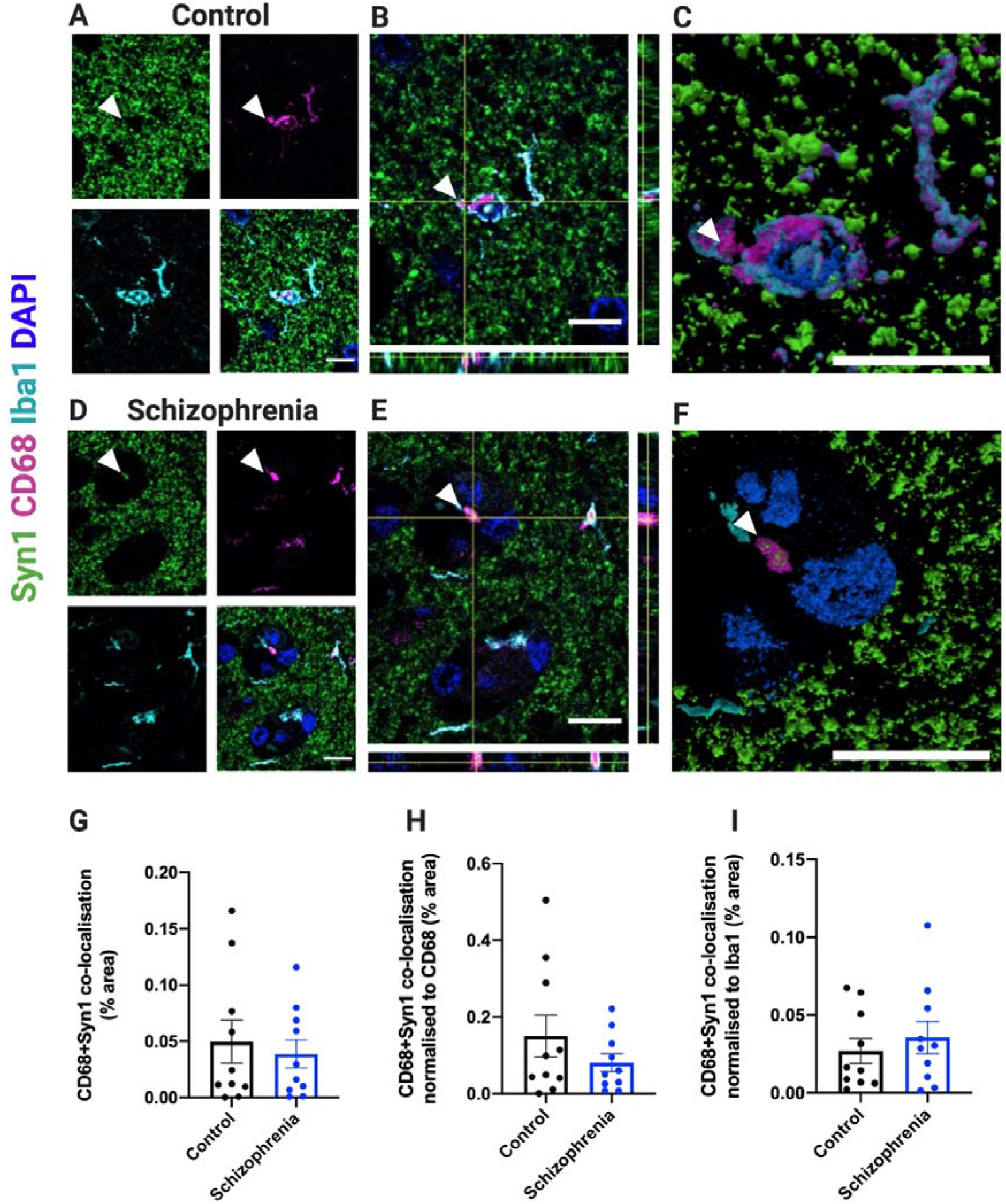
No difference found in synaptic engulfment by microglia in post-mortem tissue. Representative confocal images of the pre-synaptic marker synapsin I (green), CD68 (magenta) and Iba1 (cyan) in control (A-C) and schizophrenia tissue (D-F). Nuclei are counterstained with DAPI. A and D show individual panels of each stain and lastly the merged image, with white arrowheads pointing to sites of co-localisation between CD68 and synapsin I. B and E are expanded images of A and D, with orthogonal views indicating where CD68 and synapsin I co-localise. C and F represent 3D reconstructions from A and D, generated on ParaView. In G, the co-localisation index of CD68 and synapsin I is quantified for control and schizophrenia cases, where similar levels of synaptic engulfment by microglia are observed. By normalising each image to their respective CD68 burden or Iba1 burden there is still no statistical change in the engulfment of synapsin I by CD68. Each data point represents a mean average of 20 images taken per case, where n=10 per group. Linear mixed-effects model assessed statistical significance, considering p≤0.05 for significance. All scale bars represent 10μm.

## Discussion

In human post-mortem tissue from both patients with schizophrenia and age-matched controls, we found pre-synaptic proteins inside microglial cells in the frontal cortex of the brain, but no difference in the levels of synaptic internalisation between the two groups.

A limitation of our post-mortem tissue is that it provides a snapshot of the disease, which lacks mechanistic insight. A greater sample size in an independent cohort will be useful to assess the reproducibility of these results and to stratify by confounding variables like sex and age. This would also allow assessment of comorbid symptoms in schizophrenia, like depression or psychosis, and if such symptoms affect microglial and synaptic internalisation by the cells. However, this study is unique, as schizophrenia tissue is scarce and by the type of assessment performed.

With gliosis being reported in multiple brain disorders, we assessed microgliosis in schizophrenia. As described above, we found no differences in microglial burdens between disease and control groups. This suggests that microglial activation is not a sustained event in chronic schizophrenia, and if any changes do occur in these cells it would instead likely involve functional alterations. Previous literature looking at CD68 expression in control and schizophrenia cases has also reported a similar outcome [29]. Given that schizophrenia is not a progressive, albeit chronic disease, it is understandable that if any changes in glial dynamics were to occur, they may be seen closer to disease onset, and that by the time the brains were donated 35 years later, any changes would have subsided. This would be consistent with the observations published to visualise and quantify microglial activation *in vivo* with positron emission computed tomography (PET) using specific ligands of the translocator protein TSPO [30]. The PET studies have revealed that activated microglia are present in patients within the first 5 years of disease onset or during a psychotic state, whereas other PET studies in chronic schizophrenia have shown no difference in microglial activation between healthy controls and these patients.

Although developmental synaptic alterations, like synapse loss, have been characterised in individuals with schizophrenia [3,8], there are key unanswered questions that remain. For instance, it is not clear how the synapse elimination is mediated, the extent to which it drives behavioural symptoms, or whether it is the outcome of other disease-specific pathologies. In neurodegenerative diseases, such as Alzheimer’s disease (AD), synapse loss is a hallmark of the disease [31,32], and it has gained significant attention as it associates strongly with the cognitive decline seen in patients [33,34]. Although schizophrenia and AD have very different pathological features and the onset of the two disease is far apart, there are some common qualities that may help with understanding disease mechanisms. For example, a prominent mechanism for synaptic elimination in development is the use of the classical complement cascade (CCC), where it has been shown to sculpt neural circuits by tagging less electrically active synapses [18,20]. Recent research has now implicated complement as a signal for aberrant synapse elimination in disease [14,35]. Specifically, variants of C4 of the CCC are associated with a greater risk of developing schizophrenia [36], as well as poorer brain connectivity and schizophrenia-like behavioural deficits in mice [37].

Currently, a suggested mechanism by which complement-tagged synapses are cleared is by microglial recruitment for synaptic removal. The microglial importance in guiding cerebral circuitry has been recently described in a case study of a baby born with a homozygous mutation in the *CSF1R* gene causing a total lack of microglial cells, resulting in early death [38]. Upon autopsy, ectopic grey matter was found growing in the ventricles and impaired cell layer separation was observed in the grey matter, suggesting a faulty brain wiring due to loss of microglia. However, microglia can also be involved in abnormal synaptic elimination. Indeed, increased microglial-mediated synaptic clearance has been observed in AD mouse models [39] which was rescued in complement-deficient mice [21,40]. Of note, we have recently shown in human post-mortem tissue that in AD there is increased synaptic ingestion by microglia, and that this is exacerbated in areas near amyloid-β pathology [41]. In co-cultured neuron and microglia-like cells from human induced pluripotent stem cells from control and schizophrenia lines, increased levels of the excitatory post-synaptic protein PSD-95 was reported phagocytosed in the schizophrenia co-cultures [23]. Interestingly, this increased phagocytic activity was mainly driven by the presence of schizophrenia-derived microglia. Indeed, when schizophrenia neurons were co-cultured with microglia from control patients, the phagocytic index was reduced, indicating that in schizophrenia microglia have intrinsic differences in their phagocytic response. It is worth noting that induced stem cells are a good model for understanding human disease but represent a developmentally earlier phenotype, and not that of the age of the donor. Therefore, this supports a role for phagocytic microglia in early stages of the illness, and may explain why we did not see any changes in phagocytic ability of microglia towards synapses in chronic schizophrenia, since we are not studying the developmental time-frame.

In conclusion, here we report that microglia in human post-mortem tissue internalise pre-synaptic proteins physiologically, and that this does not appear to be altered in the chronic form of schizophrenia, at the difference to our observation in AD. Nevertheless, given the typically early onset of schizophrenia and that synapse loss is likely to have occurred years before brain collection, we cannot make assumptions on the role of microglia in synaptic clearance at the start of the disease. Looking forward, it would be interesting to study difference between young versus older cases in terms of synaptic uptake by microglia, and phenotype these changes in several brain areas to investigate any region-specific differences. Lastly, longitudinal PET imaging of the pre-synaptic marker SV2A [42–44] and TSPO microglial marker would enable exploration of any microglia-synapse association during the course of the illnesses.

## Acknowledgements

We would also like to thank our funders, specifically the UK Dementia Research Institute which receives funding from Alzheimer’s Research UK, the Alzheimer’s Society, and the Medical Research Council. We also would like to thank the Wellcome Trust for funding AJS and TLSJ. Tissue samples were obtained from The Corsellis Collection as part of the UK Brain Archive Information Network (BRAIN UK) which is funded by the Medical Research Council and Brain Tumour Research.

Authors contributed in the following ways: MT contributed in study design, performed experiments and imaging, statistical analysis, and manuscript preparation; AJS contributed in statistical analysis and manuscript editing; DB contributed by providing cut paraffin-embedded section, study design, and manuscript editing; TLSJ contributed with study design, statistical analysis, and manuscript editing. Figures created with BioRender.

---

**Supplementary Table 1:** Linear mixed-effects model outcomes from R Studio.

| Measurement | Effect size | Standard Error | p-value |
| --- | --- | --- | --- |
| Iba1 burden<br>(% area) | -0.1637 | 0.1580 | 0.315 |
| CD68 burden<br>(% area) | 0.0165 | 0.06215 | 0.794 |
| CD68+Iba1<br>colocalisation<br>(% area) | -0.0378 | 0.0792 | 0.639 |
| CD68+Syn1<br>colocalisation<br>(% area) | -0.0772 | 0.0917 | 0.413 |
| CD68+Syn1<br>colocalisation<br>/CD68 (% area) | -0.1414 | 0.0980 | 0.167 |
| CD68+Syn1<br>colocalisation<br>/Iba1 (% area) | -0.0044 | 0.0958 | 0.964 |

